# Generation of a double binary transgenic zebrafish model to study myeloid gene regulation in response to melanocyte transformation

**DOI:** 10.1101/118620

**Authors:** Amy Kenyon, Daria Gavriouchkina, Giorgio Napolitani, Vincenzo Cerundolo, Tatjana Sauka-Spengler

## Abstract

A complex network of inflammation succeeds somatic cell transformation and malignant disease. Immune cells and their associated molecules are responsible for detecting and eliminating cancer cells as they establish themselves as the precursors of a tumour. By the time a patient has a detectable solid tumour, cancer cells have escaped the initial immune response mechanisms. To date, no model exists for studying the underlying mechanisms that govern the initial phase of the immune response when transformed cells become precursors of cancer. Here we describe the development of a double binary zebrafish model designed for exploring regulatory programming of the myeloid cells as they respond to oncogenic transformed melanocytes. A hormone-inducible binary system allows for temporal control of different Ras-oncogenes (NRasK61Q, HRasG12V, KRasG12V) expression in melanocytes, enabling analysis of melanocyte transformation and melanoma initiation. This model was coupled to binary cell-specific biotagging models allowing *in vivo* biotinylation and subsequent isolation of macrophage or neutrophil nuclei for regulatory profiling of their active transcriptomes. Nuclear transcriptional profiling of neutrophils, performed for the first time as they respond to the earliest precursors of melanoma *in vivo*, revealed an intricate landscape of regulatory factors that may promote progression to melanoma including fgf1, fgf6, cathepsin H, cathepsin L, galectin 1 and galectin 3. The model presented here provides a powerful platform to study the myeloid response to the earliest precursors of melanoma.

**Summary Statement:** We present an innovative double binary zebrafish model for exploring the underlying regulatory mechanisms that govern the myeloid response mechanisms at the onset of melanoma.

## Introduction

Immune cells and their associated molecules are responsible for detecting and eliminating cancer cells and their precursors at early stages of cancer differentiation [51, 10]. A complex network of inflammation succeeds somatic cell transformation and malignant disease. Importantly, analysis of the initial response of the immune cells to cancer might lead to the discovery of targets for both prevention and treatment. As with pathogens, cancer cells can evade the immune system and as such tumour growth and ultimately the emergence of clinically detectable cancer is largely dependent on the capacity of cancerous cells to evade and manipulate the immune response [5]. Myeloid cells are equipped with sensors for damage and inflammation as well as subsequent effector mechanisms for resolving inflammation [18]. However, transformed cells are able to shape the nature of the myeloid cells and evoke an immunosuppressive response, enabling the progression of the disease [17]. Further studies are needed to understand the different inflammatory signalling pathways in tumour initiation and progression, and how they may be modulated for immunotherapy. Because we cannot predict when and where transformed cells may arise in the human host, little is known about the sterile inflammatory signalling pathways that follow somatic cell transformation closely[18].

Zebrafish is an accepted model in cancer research, with many aspects of carcinogenesis being conserved between fish and humans [50, 43]. Unlike mammalian models, the zebrafish is easily amenable to techniques that allow the study of both cancer initiation and progression[18]. Zebrafish are particularly useful in studying melanoma because their melanocytes are externally visible and large single cells can be directly visualized in living fish [8].

Previous transgenic zebrafish melanoma models used oncogenic BRAFV600 [39], NRasQ61K [13] and HRasG12V [36, 3] under the melanocyte specific microphthalmia-associated transcription factor a (mitfa) promoter, coupled to an additional mutation in the tumour suppressor gene p53. In the mitfa-driven HRasG12V transgenic line, melanoma development is rare but secondary mutations in the PI3K signalling pathway contribute to its occurrence [39, 14, 3]. Santoriello et al. subsequently developed a conditional transgenic zebrafish line, which uses the Gal4 system and the melanocyte-specific kita promoter to drive HRasG12V [48, 42]. Kita:HRasG12V induced early onset melanoma without additional mutations in tumour suppressor genes. However, some reports suggest that high levels of Gal4 expression can be toxic and may result in developmental defects [16]. Moreover, UAS sequences have been shown to be susceptible to DNA methylation leading to transcriptional silencing of the transgene, which is minimal in the first generation, but exacerbated upon propagation through later generations [25, 1].

However, the study of the myeloid response to the earliest precursors of melanoma, would require temporal control of carcinogenesis. The current zebrafish models for benign nevus and cutaneous melanoma rely on the targeted expression of a human oncogene in melanocytes, limiting the ability to control melanoma initiation. Additionally, constitutive expression of the oncogene renders maintenance of stable transgenic lines difficult because these models may develop severe tumours before the fish reach reproductive age [37].

To overcome these limitations we set out to develop a inducible system of melanoma initiation. Here we report the generation of a novel inducible zebrafish model, specifically designed to study the onset of melanoma using the mifepristone-inducible LexPR system developed by Emelyanov and Parinov [16]. Additionally, we have developed an innovative transgenic zebrafish model system for *in vivo* biotinylation and subsequent isolation of neutrophil (previously described in [28]) and macrophage nuclei based on the Biotagging method [Trinh et al.]. Using this approach, it is possible to perform genome-wide analysis of the active nuclear transcriptomes of neutrophils and macrophages. By combining the Biotagging approach that enables isolation and analysis of either neutrophil or macrophage nuclei with the inducible model for melanoma initiation, we have created a powerful double binary system for regulatory profiling of myeloid cells that respond to the precursors of melanoma. In a proof on concept analysis, we report for the first time that neutrophils responding to the melanoma initiation upregulate a number of factors which may promote melanoma progression.

## Results

### Generation of a binary inducible melanoma model

To generate an inducible model for the initiation of melanoma tumorigenesis, we adapted a hormone-inducible binary system for targeted gene expression in zebrafish, previously used to drive liver carcinogenesis [16, 37]. In this system the transcriptional activator expressed by the transgenic driver is a ligand-dependent chimeric transcription factor, termed LexPR transactivator. LexPR transactivator is a fusion protein containing the DNA-binding domain of the bacterial LexA repressor, a truncated ligand-binding domain of the human progesterone receptor and the activation domain of NF-κB/p65 protein. The effector transgenic fish line harbours the operator-promoter sequence, consisting of the synthetic LexA operator fused to a minimal 35S promoter sequence from the Cauliflower mosaic virus, upstream of the reporter gene. In the presence of the progesterone agonist, mifepristone, that binds to its ligand-binding domain within the LexPR transactivator, LexPR activates the transcription of the target reporter gene by binding to the LexA binding sites within the LexA operon (LexOP) positioned upstream (Figure 1A).

**Figure 1.**
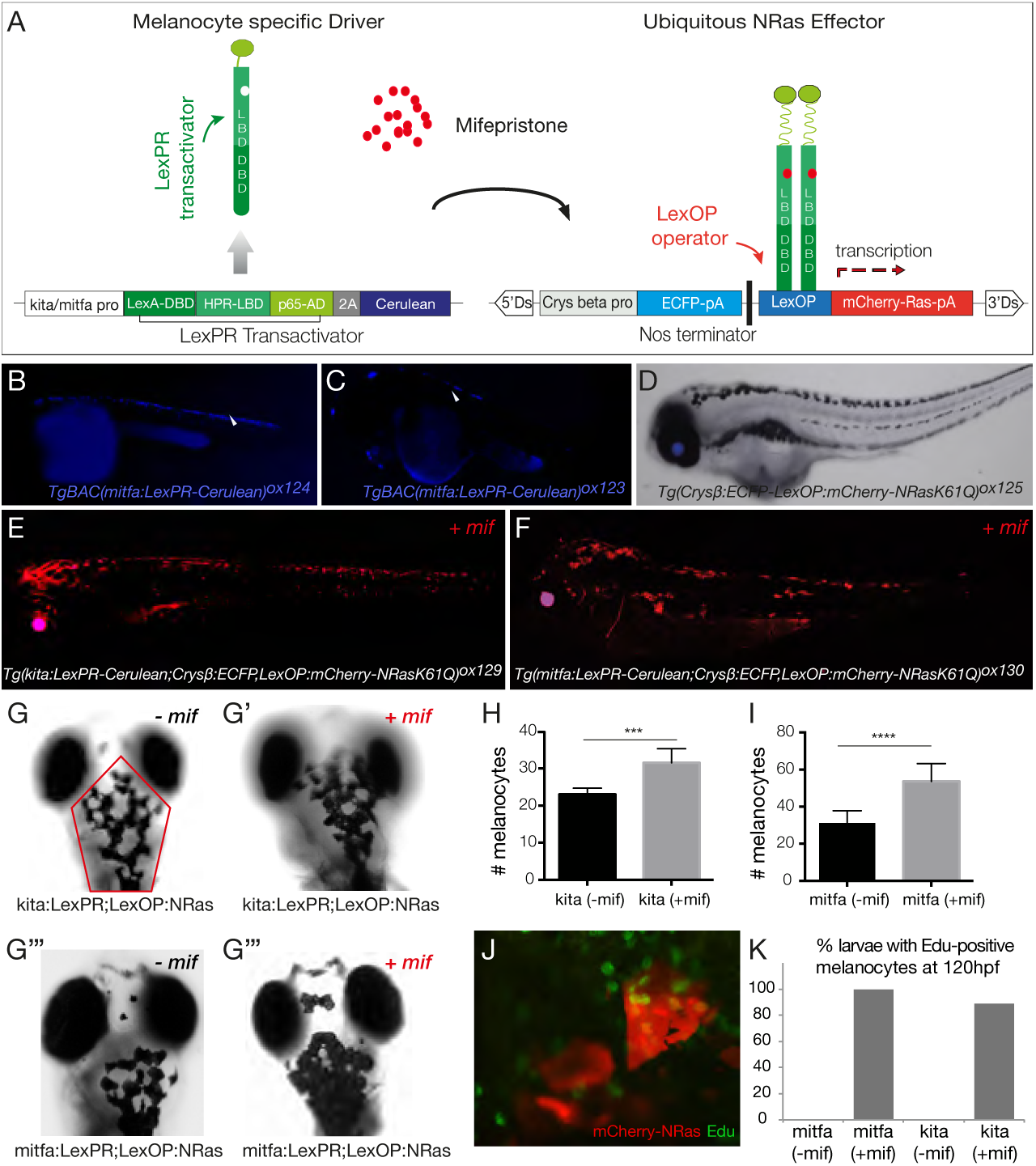
LexPR mifepristone-inducible model for melanocyte transformation(A) Diagram of LexPR/LexOP inducible system. The LexPR driver cassette consists of LexPR transactivator and a Cerulean reporter under the control of melanocyte-specific promoters, kita or mitfa. The effector cassette contains the Lex operator sequence (LexOP) fused to the mCherry-Ras oncogene. mCherry-Ras fusion is transcribed in trans only in the presence of mifepristone, when the LexPR transactivator-mifepristone complex binds to the LexOP sequence upstream of the oncogene expression cassette. **(B-C)** mitfa:LexPR-Cerulean (*TgBAC(mitfa:LexPR-Cerulean)*]^*ox124*^) and kita:LexPR-Cerulean (*TgBAC(kita:LexPR-Cerulean)*^*ox123*^) transactivator driver lines are characterised by expression of LexPR transactivator and its cognate fluorescent protein Cerulean specifically in melanocytes. **(D)** LexOP:mCherry-NRas effector line (*Tg(Crys β: *ECFP-LexOP:mCherry-NRasK61Q)*^ox125^*) shows Cerulean expression in the eye, but no oncogene transcription. **(E-F)** The melanocyte-specific expression of mCherry-NRas in transgenic larvae harbouring both a transactivation driver and an oncogene effector alleles (*Tg(kita/mitfa:LexPR-Cerulean;LexOP:mCherry-NrasK61Q)*^ox129/ox130^) is activated by addition of 1 *μ*M mifepristone to the embryo rearing solution. (G-G’) Dorsal views of the head of 5dpf *Tg(kita:LexPR-Cerulean;LexOP:mCherry-NRas)*^ox129^ larvae without **(G)** and with **(G’)** mifepristone-dependent oncogene activation. **(G”-G”’)** Dorsal views of the head of 5 dpf *Tg(mitfa:LexPR-Cerulean;LexOP:mCherry-NRask61Q)*^ox130^ larvae without **(G”)** and with **(G”’)** mifepristone-dependent oncogene activation. **(H-I)** Quantification of the number of melanocytes in the region indicated by the red outline in **G**, comparing larvae from kita:LexPR;LexOP:NRasK61Q (n=8) **(H)** and mitfa:LexPR;LexOP:NRasK61Q (n=9)**(I)** model, with and without oncogene activation (± mif). Statistical significance was determined by two-tailed unpaired students t-test with Welch’s correction. **(J)** Proliferative activity of transformed melanocytes (red) shown by Edu incorporation (green). Embryos were pulsed with Edu by pericardial injections at 96 hpf, when all melanocytes are already postmitotic. NRas-transformed melanocytes show continuous proliferation. **(K)** Quantification of number of kita:mCherry-NRasK61Q and mitfa:mCherry-NRasK61Q larvae with proliferating (Edu-positive) melanocytes, when reared in the presence (+ mif) or absence (-mif) of mifepristone (n=6-8).

In our system we placed the LexPR transactivator, followed by Cerulean reporter, under the control of the melanocyte-specific promoters kita and mitfa using BAC-mediated transgenesis (Fig. S1). Two melanocyte-specific driver lines (*TgBAC*(*kita:LexPR-Cerulean*)*^ox123^* (heron referred to as kita:LexPR-Cerulean) and *TgBAC*(*mitfa:LexPR-Cerulean*)*^ox124^* (heron referred to as mitfa:LexPR-Cerulean) were generated (Figure 1 B-C).

To generate the effector lines, different Ras-oncogenes (NRasK61Q, HRasG12V and KRasG12V) fused to a mCherry fluorescent reporter were cloned downstream of LexOP operator. The effector constructs were placed within a non-autonomous maize dissociation (Ds) element, which allows for effective activator (Ac)-mediated transposition using the transposase Activator (Ac)/ Dissociation (Ds) (AcDs) system [16] (Fig. S1). This system is extremely efficient and results in a very high number of independent integrations into the zebrafish genome, thus allowing for high level expression of the Ras effectors activated by LexPR in the context of oncogenic transformation. Three Ras-effector lines were generated: *Tg*(*Crysβ:ECFP-LexOP:mCherry-NRasK61Q*)^ox125^ (hereon referred to as LexOP:mCherry-NRasK61Q), *Tg*(*Crysβ:ECFP-LexOP: mCherry-HrasG12V*)^ox126^ and *Tg*(*Crysβ:ECFP-LexOP:mCherry-KrasG12V*)^ox127^. The Ras-effector lines all contain the selection marker ECFP under the control of lens-specific zebrafish crys β promoter (crystallin β), which results in blue fluorescence in the lens. This can be used to screen for transgene integration, without driving the oncogene (Figure 1D). When a driver line is crossed to an effector line, in the presence of the synthetic progesterone ligand, mifepristone, the mCherry-Ras oncogene is transcribed only in melanocytes. Two different melanocyte-specific driver lines (kita and mitfa) and three different mCherry-Ras oncogene lines were generated in this study paving the way for multiple combinations of the system. We characterised two of those lines, *Tg*(*kita:LexPR-Cerulean; Crys β:ECFP-LexOP:mCherry-NRasK61Q*)*^ox129^* (Figure 1E) and *Tg*(*mitfa:LexPR-Cerulean; Crys β:ECFP-LexOP:mCherry-NRasK61Q*)*^ox130^* (Figure 1F) (hereon referred to as kita:LexPR;LexOP:NRasK61Q and mitfa:LexPR;LexOP:NRasK61Q, respectively) and showed that when embryos positive for both alleles are reared in the presence of mifepristone, melanocyte-specific expression of the mCherry-NRas fusion protein is detected. Transformed melanocytes are seen visible in pairs or clumps of transformed red cells as shown by the white arrows, indicative of their proliferative potential (Figure 1 E-F). In the mitfa:mCherry-NRasK61Q genotype, low levels of melanin were sometimes observed in melanocytes expressing mCherry (data not shown). To quantify the degree transformed melanocytes proliferation in kita:mCherry-NRasK61Q and mitfa:mCherry-NRasK61Q embryos, we counted the total number of melanocytes within the outlined area in the cranial region in the presence and absence of mifepristone (± mif), (Figure 1G-G”’). Both transgenic lines showed a statistically significant increase in the number of melanocytes in the region of interest (Figure 1H-I).

To assay for proliferative activity of transformed melanocytes, the kita:mCherry-NRasK61Q and mitfa:mCherry-NRasK61Q embryos were pulsed with EdU using pericardial injection at 96 hours post fertilization (hpf). In zebrafish, all melanocytes are post-mitotic by 60 hpf, when the larval pigmentation pattern is largely complete [27, 26]. Embryos were allowed to develop until the following day, when they were fixed and stained with EdU Click-iT reaction mixture (green), together with RFP-booster to amplify the mCherry signal labelling the transformed melanocytes. Consistent with melanocyte proliferation at 96 hpf, we show that Edu incorporates into the melanocytes (Figure 1 J) of kita:mCherry-NRasK61Q and mitfa:mCherry-NRasK61Q embryos, reared in presence of mifepristone (+ mif). We quantified this effect by assessing the number of embryos in which we detect at least 60% of visible melanocytes that have incorporated Edu. We found that 89% and 100% of the kita:mCherry-NRasK61Q and mitfa:mCherry-NRasK61Q embryos respectively, harbour continuously proliferating transformed melanocytes. This was in sharp contrast to stage- and genotype-matched control embryos, reared in the absence of mifepristone (-mif), where Edu was not detected in melanocytes in any instance (Figure 1 K).

### Establishment of a binary system to carry out regulatory profiling in myeloid cells

Genome-wide analysis of the regulatory networks that govern the inflammatory response *in vivo* is complicated by the fact that relatively small numbers of responding cells are present within complex tissues. The current zebrafish lines for studying macrophage and neutrophil function make use of fluorescent reporters under the control of the mpeg1 and mpx promoters respectively [15, 40]. These lines have largely been used for functional studies at the cellular level, allowing for a detailed examination of host-pathogen interactions, comparative analysis of macrophage and neutrophil behaviours responding to inflammation, and the dynamic interaction between the two cell types [15, 40]. Based on the Biotagging method [Trinh et al.], we have developed a zebrafish model to study the gene regulatory landscape of macrophages and neutrophils by cell specific *in vivo* biotagging of nuclei, thereby enabling us to obtain the active transcriptome of these cell types (Figure 2A). In our system, the Biotagging driver line harbours cell-specific (macrophage or neutrophil) expressed *E. coli* biotin ligase BirA. Myeloid BirA driver lines *TgBAC*(*mpeg1:BirA-Citrine*)^ox122^ (hereon referred to as mpeg1:BirA-Citrine (Figure 2B) and *TgBAC*(*mpx:BirA-Citrine*)*^ox121^* (hereon referred to as mpx:BirA-Citrine (Figure 2C) (described in [28]) were generated using BAC-mediated transgenesis as previously described [Trinh et al.](Figure S1). Previously characterised effector line *Tg*(*βactin:Avi-Cerulean-RanGap*)^ct700a^ (hereon referred to as bactin:Avi-Cerulean-Rangap) [Trinh et al.] features ubiquitous expression of a zebrafish-compatible version of Avi-tagged nuclear envelope-associated fusion protein. In addition to the biotinylatable Avi-tag, this effector protein consists of the carboxyl-terminal domain of Ran GTPase-activating protein1, RanGap1, which acts as an outer nuclear envelope-targeting sequence, and a fluorescent variant Cerulean allowing visualisation of the fusion protein (Figure 2D). When the driver line is crossed to the effector line, in the specific cells (neutrophils or macrophages) that carry both Biotagging alleles (driver and effector) the Avi-tagged fusion protein gets biotinylated and localizes to the outer nuclear envelope. This process effectively results in the biotinylation of nuclei in a cell-specific (macrophage or neutrophil) fashion, allowing isolation of nuclei from these cell types using streptavidin-coated magnetic beads. Subsequently, the content of these nuclei can be analysed in a genome-wide fashion by Next Generation Sequencing (NGS) techniques. The use of mpeg1:BirA-Citrine and mpx:BirA-Citrine drivers combined with the Avi-Cerulean-Rangap effector offers the unique possibility to interrogate initial, often crucial regulatory responses in specific cell types. Given that cell-specific biotinylation onsets very shortly after the cells start expressing BirA, there’s no time lag in the analysis, as is the case for cells isolated based on fluorescent reporters which require a certain a amount of time to mature. This is particularly crucial for fast developing zebrafish embryos.

**Figure 2.**
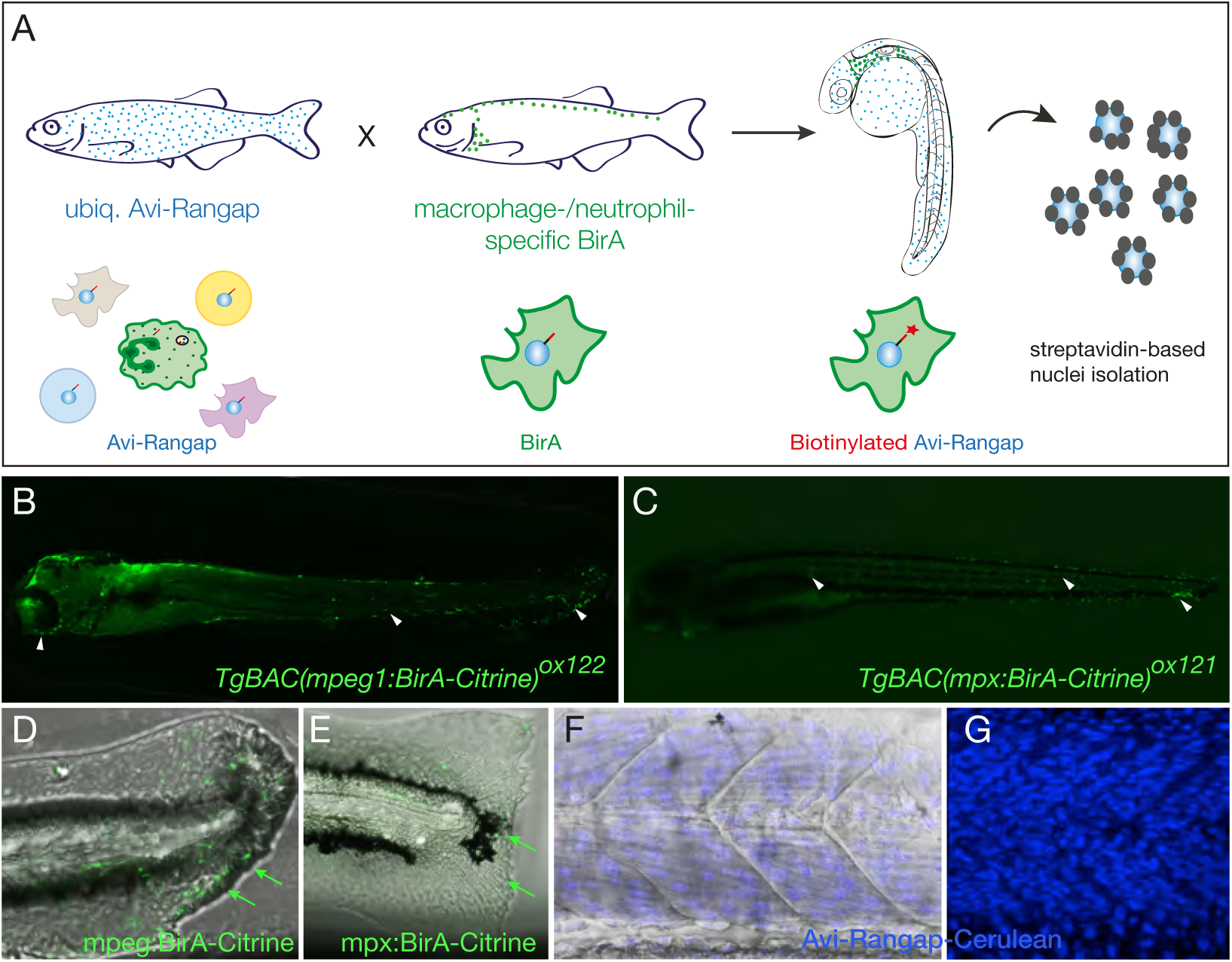
Binary transgenic zebrafish model for regulatory profiling of myeloid cells(A) Schematic of myeloid nuclear biotagging system. When biotagging effector transgenic zebrafish line ubiquitously expressing Avi-tagged Rangap for biotinylation of nuclei is crossed to Biotagging driver line expressing BirA in myeloid-specific manner, only the macrophage or neutrophil nuclei will be biotinylated. **(B)** Transgene expression in macrophage-specific BirA driver, *TgBAC(mpeg1:BirA-Citrine)*^ox122^, is amplified with αGFP-Alexa488 to show expression in macrophages as indicated by the white arrowheads. **(C)** *TgBAC(mpx:BirA-Citrine)*^ox121^ biotagging transgenic driver shows neutrophil-specific expression as indicated by the white arrows. **(D-E)** Tailfins of transgenic fish transected at 3dpf, show responding macrophages in mpeg1:BirA-Citrine **(D)** and neutrophils mpx:BirA-Citrine **(E)** larvae at 5 hours post injury (5hpi), as indicated by green arrows. **(F-G)** Projection of confocal microscope images of Tg(bactin:Avi-Cerulean-Rangap)^ct700a^ show ubiquitously expressing biotagging effector specifically on nuclei, across somite region of the embryo, with **(F)** and without **(G)** bright field background. The images in F and G are taken in different embryos.

To show that the mpeg1/mpx driver lines generated in this study faithfully recapitulate larval inflammatory phenotypes, tailfin transections [40] were carried out at 3 days post fertilization (dpf) in mpeg1:BirA-Citrine and mpx:BirA-Citrine embryos, respectively. Injured specimens were fixed at 5 hours post injury (hpi) and stained using anti-GFP antibody to amplify the Citrine signal and thus detect macrophages and neutrophils. Convincingly, both mpeg1:BirA-Citrine (Figure 2D) and mpx:BirA-Citrine (Figure 2E) showed infiltration of macrophages and neutrophils in the site of injury.

### Generation of a double binary system to study the myeloid response to transformed melanocytes

To study the initial myeloid response to oncogene-transformed melanocytes, we have generated a double binary system in zebrafish, by coupling the inducible LexPR driver/effector to the Biotagging system for nuclear transcriptional profiling of macrophages and neutrophils. In the simplest form, kita/mitfa:LexPR-Cerulean (Figure 3, fish a) is crossed to crys *β*: ECFP-LexOP:mCherry-NRas/HRas/KRas (Figure 3, fish b) to generate inducible transgenic fish carrying both alleles (Figure 3: Binary System 1, fish ab). Similarly, mpx/mpeg1:BirA-Citrine (Figure 3 fish c) was crossed to *β* actin:Avi-Cerulean-Rangap (Figure 3, fish d) to generate mpx/mpeg1:BirA;bactin:Avi-Rangap (Figure 3: Binary System 2, Fish cd). Subsequent analyses can then be carried out in embryos by crossing binary system 1 (Figure 3, fish ab) with binary system 2 (Figure 3, fish cd).

**Figure 3.**
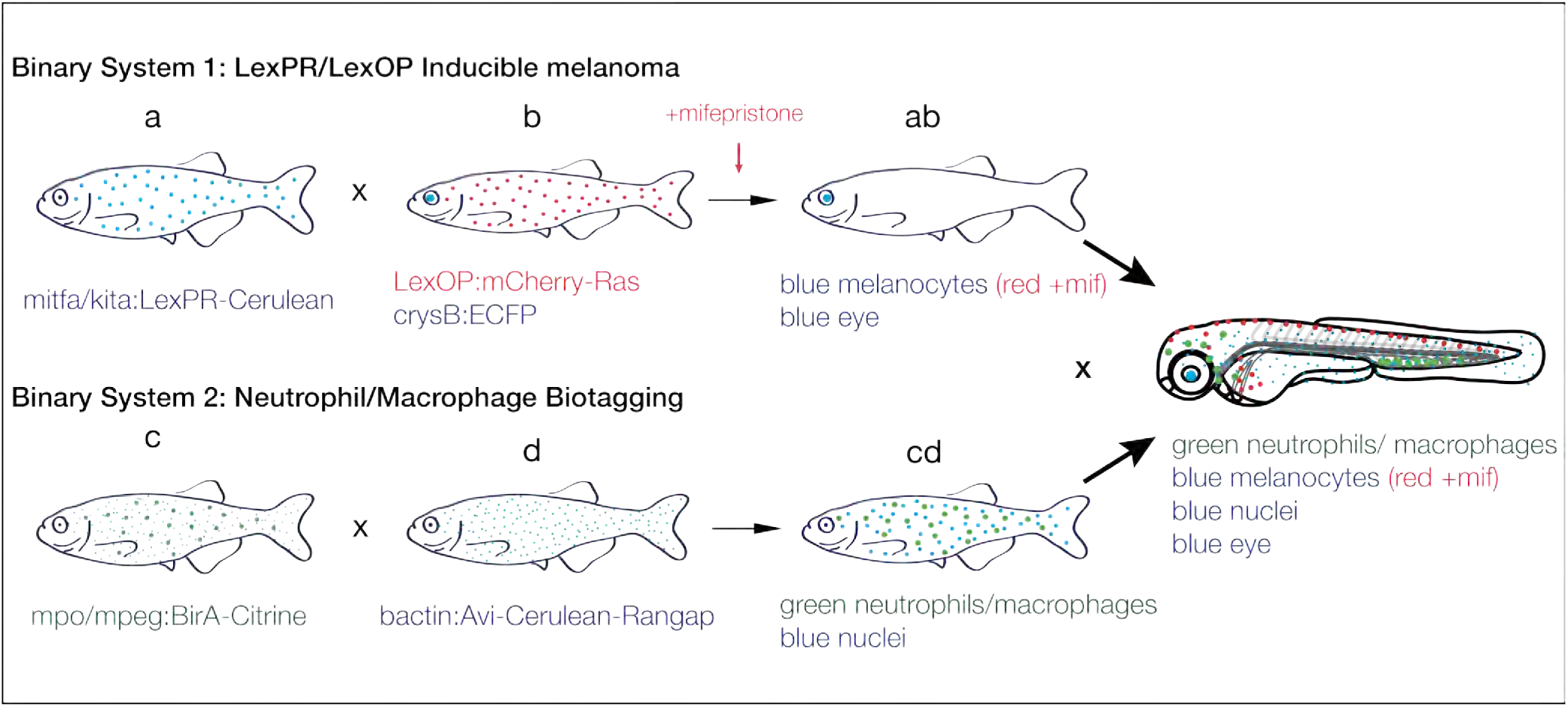
Schematic representation of double binary system to study the myeloid response to melanoma. **Binary System 1:** The LexPR transactivator expressed under the control of a melanocyte-specific promoter activates in trans the mCherry-Ras oncogene in the presence of a specific ligand (mifepristone). Fish a) Melanocyte-specific LexPR transactivator with Cerulean reporter. Fish b) Lex Operon controlled mCherry-Ras with crystallin-specific ECFP expression. **Binary System 2:** Nuclear envelope protein tagged with a biotin acceptor peptide (Avi-tag) is biotinylated in macrophages/neutrophils allowing for cell-specific isolation of nuclei and genome-wide analysis. Fish c) Macrophage/neutrophil-specific biotin ligase, BirA, with Citrine reporter. Fish d. Ubiquitous Avi-tagged nuclear envelope protein with Cerulean reporter. Fish a crossed to Fish b results in Fish ab (blue eyes and blue melanocytes, red in the presence of mifepristone). Fish c crossed to Fish d in Fish cd (green macrophages/neutrophils and blue nuclei). Fish ab crossed to Fish cd yields embryos to be analysed (green neutrophils/macrophages, blue melanocytes or red in the presence of mifepristone, blue nuclei, blue eyes).

### Kita-driven oncogenic NRas melanocytes are immunogenic

Since NRasK61Q is the most relevant oncogene to human melanoma with 30% of melanomas having contributing mutations in NRas [8], it was prioritized for the purpose of this study. To show that NRasK61Q-transformed melanocytes generated by the inducible LexPR binary system were immunogenic, kita:LexPR;LexOP:NRasK61Q fish were in-crossed and embryos were grown to 24 hpf, when mifepristone was added to the water. Larvae with NRasK61Q-transformed melanocytes were reared until 5 dpf, fixed, and stained for mpeg1 and mpx antibodies to detect macrophages and neutrophils, respectively. Macrophages were seen in the vicinity as well as directly interacting with the NRasK61Q-transformed cells (Figure 4A), in the zebrafish larvae. The neutrophils appeared to form a ring-like structure surrounding the transformed melanocytes (Figure 4B).

**Figure 4.**
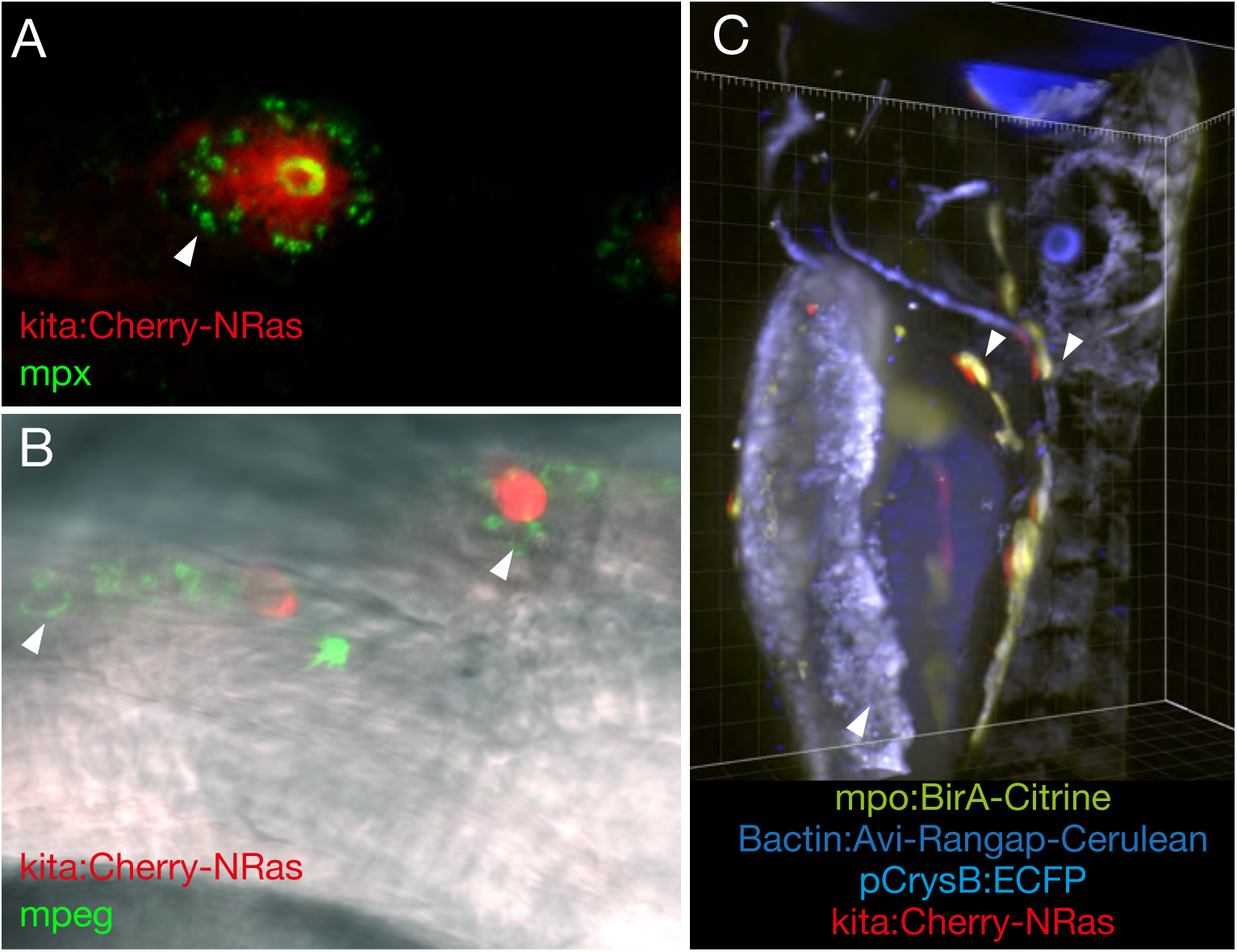
mCherry-NRasK61Q transformed melanocytes are immunogenic (A-B) The melanocyte-specific expression of mCherry-NRas in *Tg(kita;LexPR-Cerulean;pCrysβ:ECFP-LexOP:mCherry-NRasK61Q)*^ox129^ was activated by addition of mifepristone to the embryo rearing solution at 24 hpf. At 5 dpi, confocal images show macrophages (green) and neutrophils (green), detected with mpeg1 and mpx antibodies respectively, interacting with a mCherry-NRas-transformed melanocyte (red) as indicated by the white arrows. **(C)** kita:LexPR;LexOP:NRasK61Q were crossed to mpx:BirA;bactin:Avi-Rangap, and mCherry-NRas activated by addition of mifepristone. 5dpi larvae harbouring all four (blue eye, blue nuclei, green neutrophils and red melanocytes) shows neutrophils in close proximity to transformed melanocytes as captured on a Zeiss Z1 lightsheet microscope and indicated by the white arrows.

Similarly, melanoma-inducing binary system 1 transgenic line, kita:LexPR;LexOP:NRasK61Q was crossed to neutrophil Biotagging binary system 2 transgenic, mpx:BirA;bactin:Avi-Rangap (*Tg*(*mpx:BirA-Citrine;bactin:Avi-Cerulean-Rangap*)^ox128^). Oncogene expression was activated in the melanocytes and goblet cells by addition of mifepristone to the water at 24 hpf. Larvae were fixed and imaged at 5 dpf. In a fish harbouring all four alleles which features a blue eye, blue nuclei, green neutrophils and NRasK61Q-transformed red melanocytes/goblet cells, the green neutrophils were localised in the vicinity of the transformed melanocytes (Figure 4C).

### Transcriptional analysis of neutrophils in response to NRasK61Q-transformed melanocytes by ***in vivo*** biotagging

To test the efficacy of this model, we carried out a proof of concept study and performed transcriptional profiling of neutrophils responding to transformed melanocytes, with the aim to identify components of the cellular response which might contribute to tumour progression. All subsequent experiments were carried out by crossing kita:LexPR;LexOP:NRasK61Q inducible melanoma system for NRasK61Q-transformation of melanocytes to mpx:BirA;bactin:Avi-Rangap biotagging system allowing to isolate neutrophil nuclei responding to oncogenic transformation. NRasK61Q expression in the melanocytes of resulting embryos was driven by addition of mifepristone to the water at 24 hpf. Embryos expressing mCherry-NrasK61Q in the melanocytes were selected for the profiling experiments, allowed to develop until 5 dpf and used for analysis. Approximately 45 larvae from each experimental (+mif) and control (-mif) condition were used per experiment. A total of 3 biological replicates were collected. In each experiment, prior to nuclei isolation and analysis, cranial regions containing major foci of melanocyte oncogenic transformation were dissected to eliminate anterior yolk sac and Intermediate Cell Mass (ICM), thus reducing the background levels from non-responding neutrophils (Figure 5A) [4]. By focussing our analysis on the isolated transformed mCherry-expressing regions in the head, we enriched the samples in the environing neutrophils responding to transformed melanocytes. Moreover, mCherry-NRas expressing goblet cells found in the trunk region were excluded from the analysis.

**Figure 5.**
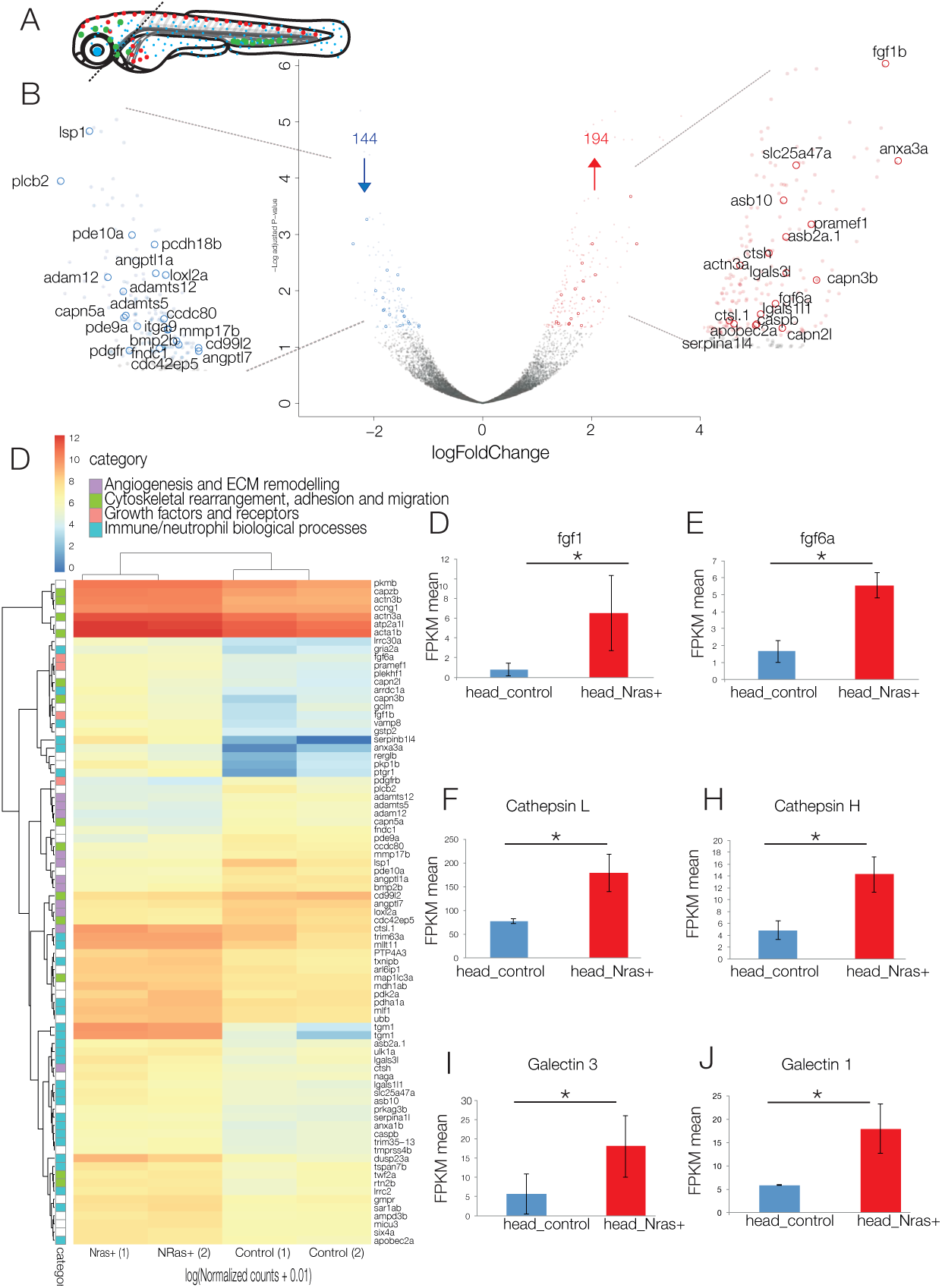
Analysis of technical replicates to obtain differentially expressed genesA) Schematic of dissection of 5 dpf larvae for nuclear profiling of neutrophils in the presence or absence of NRask61Q transformed cranial melanocytes. Cranial regions are dissected away from the trunk to eliminate neutrophils from the anterior yolk sac and ICM, contaminating sources of non-responding neutrophils. **(B)** Volcano plot of differential expression analysis of activated NRasK61Q and controls embryos shows the relationship between p-value and log fold change (red-upregulated; blue-downregulated) show 194 upregulated genes (red) and 144 downregulated genes (blue) in cranial neutrophils. **(C)** Heatmap shows the log10 (normalised counts (NMCT) +0.01) of selected differentially expressed transcripts (adjusted p-value ί 0.05). Red - high expression. Yellow - medium expression. Blue - low expression. Selected differentially expressed transcripts are further classified into subcategories of Angiogenesis and ECM remodelling (purple), cytoskeletal rearrangement, adhesion and migration (green), growth factors and receptors (pink) and immune/neutrophil biological processes (turquoise). **(D-I)** Fragments per kilobase of transcript per million mapped reads (FPKM) are represented graphically to show statistically significant upregulation in controls vs activated NRasK61Q samples for fgf1, fgf6a, cathepsin H, cathepsin L, galectin 3 and galectin 1.

’Biotagged’ nuclei were isolated from dissected tissue by nuclei pulldown [Trinh et al.]. Nuclear RNA pools were extracted and libraries prepared using SMART-seqTMv4 technology for cDNA synthesis and amplification for small cell numbers followed by Illumina Nextera library preparation. Libraries were sequenced on a Next Generation Sequencing (NGS) platform.

Scatterplots of technical replicates show reproducibility between the 3 independent experiments (Figure S2). Differential expression analysis comparing active transcription in neutrophil nuclei of embryos with Nras transformed melanocytes versus control embryos identified 194 upregulated genes and 144 downregulated genes with a statistical significance cut-off of p¡0.05 (Figure 5B-C, Supplementary data).

The analysis of RNA-seq datasets showed that a number of genes which may contribute to melanoma progression were upregulated in neutrophils upon engaging with transformed melanocytes. Those include mediators of tumour growth, angiogenesis and ECM remodelling (Figure 5D), suggesting their potential pro-tumorigenic role in promoting progression to melanoma. In addition, we found significant enrichment for a number of factors involved in neutrophil migration, adhesion and cytoskeletal rearrangement as well as key factors involved in neutrophil biological processes. Of particular interest to us were factors that may contribute to malignant progression. We detected a significantly significant increase in members of the fibroblast growth factor (FGF) family, fgf1 anf fgf6 (Figure 5E-F), known to control cellular proliferation, survival, migration and differentiation. Uncontrolled FGF signalling, implicated in diverse tumour types can drive tumorigenesis [47]. Notably we detect a statistically significant increase in cathepsin H and cathepsin L (Figure 5G-H). Cathepsins are a group of lysosomal proteinases or endopeptidases, of which different members can play a role in different tumorigenic processes including proliferation, angiogenesis and metastasis [45]. Furthermore, we see a statistically significant upregulation in members of the galectin family, galectin 3 and galectin 1 (Figure 5 I-J), a group of proteins that bind β-galactoside whose expression correlates with tumour development and invasiveness [9].

## Discussion

Although progress has been made in understanding melanoma biology, the incidence of melanoma continues to increase and the prognosis for melanoma patients still remains poor [3]. Here we report a novel inducible model to study melanoma initiation that allows for the spatio-temporal control of oncogene expression in zebrafish. Additionally, we have developed a binary model designed for regulatory nuclear profiling of myeloid cells. Coupled together these systems provide the potential to understand the mechanisms by which macrophages and neutrophils may promote melanoma progression or conversely eliminate cancer cells as they establish themselves as the precursors of a tumour. Importantly, in combination or as individual systems, both binary models provide the potential to study different aspects of melanoma biology, including the transformation of melanocytes, myeloid immunity and immunotherapy. For example, the larval phenotype observed in this zebrafish melanoma model can provide a powerful platform for testing drug targets for pre-clinical testing of therapeutics. Because of the larval phenotype that we see, the mCherry positive transformed cells can provide a direct biological readout in fast, easy to score chemical screens, aimed at finding compounds or drugs that may revert the overproliferation phenotype observed in melanocytes. Previous studies have shown that activated components of the Ras signalling pathway drive a tumour-promoting inflammatory response [18]. However, these findings do not address the difference in the inflammatory response that different driver mutations may elicit. In our model, the same driver line can be crossed to multiple oncogene effector lines. This provides further opportunities to study mechanistic detail of melanoma initiation and progression with different driver mutations.

In this proof-of-concept study, we chose to use this model system to profile the neutrophil response to NRasK61Q transformed melanocytes using the kita driver. Neoplastic cells may only initiate tumour formation in the context of a supportive tumour microenvironment (TME), largely influenced by myeloid cells. Although an immunosuppressive mechanism for neutrophils has been accepted in solid tumours, the role for neutrophils in tumour initiation remains to be elucidated. Although it is now accepted that tumour-associated neutrophils (TANs) develop a pro-tumourigenic phenotype, largely driven by the presence of TGF-*β*, inhibition of TGF-*β*; can modulate the cells to be tumouridical [21]. In untreated tumours, neutrophils were reported to contribute to tumourigenesis by secreting factors that promote tumour growth, ECM remodelling, and suppress the immune system [23]. We show here for first time that neutrophils responding to early melanoma onset may provide an early source of fibroblast growth factors (FGFs), fgf1 and fgf6. Given the multiple functional roles for FGF signalling in different tumours, the strong enrichment in FGF components in the neutrophils responding to oncogene-transformed melanocytes suggests a pro-tumorigenic role in promoting progression to melanoma. For example, uncontrolled fibroblast growth factor signalling in tumours can lead to both expansive tumour growth and progression to metastasis [29]. Furthermore, as FGF signalling can also promote angiogenesis, it is possible that FGF early role in melanoma may be linked to angiogenesis, which not only promotes tumour growth, but, also the advancement from a pre-malignant to a malignant phenotype [52, 34]. In line with this interpretation, excessive angiogenesis, a hallmark of melanoma, and the progression from a radial growth phase to a vertical growth phase, has been shown to require high angiogenic activity dependent on FGF1, FGF2 and VEGFA [34]. In particular, FGF1 is a direct activator of phosphatidyl inositol 3-kinase (PI3K)-AKT signalling in endothelial cells known to initiate migration and invasion [52]. These findings are particularly meaningful given that naturally targeting paracrine signalling in the form of growth and angiogenic factors is poised to provide exciting targets for cancer therapy.

Traditionally, cysteine cathepsin proteases are largely accepted as degradative enzymes of the lysosome. More recently, secreted cathepsins have emerged as potent effectors of multiple processes during tumour development including the turnover and degradation of the ECM as well as processing or degradation of various growth factors, cytokines and chemokines. Consequently, cathepsins may promote tumour growth, tissue invasion and metastasis (Olson and Joyce, 2015). In a model of pancreatic islet carcinogenesis, deletion of Cathepsin H significantly altered angiogenic switching of pre-malignant hyperplastic islets and ultimately a reduction in the number of tumours formed [24]. Cathepsin H activity was largely attributed to macrophages in close proximity to the blood vessels [24]. Although Cathepsin L activity has been demonstrated in various tumour types, the mechanism appears to be context dependent [6]. Expression of Cathepsin L in highly metastatic B16F10 murine melanoma cells contributed to melanoma cell invasion and migration [52]. Here we provide evidence for neutrophils as a source of Cathepsin H and Cathepsin L in melanoma initiation.

Another interesting class of molecules implicated in tumour progression are the galectins, a family of lectin carbohydrate binding proteins, that function both intracellularly and extraceullularly [20]. Multiple examples exist in the literature for the role of galectins in tumour progression including facilitating neoplastic transformation, tumour cell survival, angiogenesis and tumour metastasis [32]. Galectin 1, for example, increases cellular growth and exerts its effects by binding to the ECM as well as other cells [32]. In melanoma, Galectin 1 has been thought to cause resistance of melanoma cells to cytotoxic stimuli and to enhance angiogenesis [35]. Extracellular Galectin 3 may influence tumour progression by impeding the endocytosis of key receptors including TGF*β* and EGF while at the same time inducing endocytosis of *β* 1 integrins [22, 38]. Extracellular Galectin 3 secreted by tumour cells has also been shown to induce angiogenesis [32]. Furthermore, since galectins are expressed by immune cells, both intracellular and extracellular galectins may serve to regulate immune cell function [32]. Galectin 1 inhibits the release of inflammatory mediators by neutrophils and reduces transendothelial migration of the cells in response to inflammatory stimuli [32, 30]. Therefore, targeting galectins derived from neutrophils in benign neoplasms may serve to prevent malignant transformation by promoting a pro-inflammatory environment. Galectin 3, on the other hand, can function as a chemokine attracting both monocytes and macrophages [41]. In this capacity Galectin 3 may facilitate tumour progression driven by macrophages.

The zebrafish embryo has previously been used as a model organism to study the interactions between macrophages and neutrophils and oncogene-transformed melanocytes and goblet cells using a Kita:HRasG12V transgenic line [19]. This study revealed a recruitment of both neutrophils and macrophages to oncogene-transformed cells. The authors reported frequent cytoplasmic tethers between the immune cells and transformed cells, including neutrophils, as well as phagocytosis of transformed cells by macrophages ([19]. In agreement with this data, we show, at transcriptional level, an upregulation in signalling molecules involved in cell migration as well as cytoskeletal reorganisation in neutrophils.

In conclusion, we developed a the double binary transgenic system for studying the myeloid response to the earliest precursors of melanoma and demonstrated that is a powerful resource, offering exciting potential to reveal novel inflammatory mediators that may contribute to tumour promotion and progression. This unbiased, genome wide approach, carried out at the earliest stages of somatic cell transformation *in vivo*, is currently the only study of its kind. The ensuing gene set provides a list of interesting, novel pathways and specific candidates to explore further in human melanoma biology. The results described here contribute to the growing body of evidence that suggests a tumour-promoting role of neutrophils in cancer biology and provides insight into the mechanisms by which neutrophils can be harnessed for immunotherapy.

## Materials and Methods

### Zebrafish Maintenance and Strains

This study was carried out in compliance with local ethical approval from the University of Oxford and using procedures authorized by the UK Home Office in accordance with UK law (Animals Scientific Procedures Act 1986). Zebrafish were maintained as described [49]. Wild-type embryos for transgenesis were obtained from AB or AB/TL mix strains.

### Transgenic fish line generation and maintenance

pDs(cry:ECFP-LexOP:Cherry) construct was a kind gift from Dr Alexander Emelyanov and Dr Sergey Parinov [16]. pCrys *β*: ECFP-LexOP:mCherry-NRasK61Q, pCrys *β*: ECFP-LexOP:mCherry-HrasG12V and Crys *β*: ECFP-LexOP:Cherry-KrasG12V were cloned by Infusion cloning using pBabe-NRasK61 (plasmid #12543; Addgene), mEGFP-HRasG12V (plasmid #18666; Addgene) and pEFm.6 HA-KRasv12 (a gift from Prof Xin Lu) as donor vectors. The optimised ORF of LexPR transactivator, which eliminates all putative donor and acceptor sites that could interfere with proper production of the LexPR transcript, was synthesised based on the coding sequence obtained from Dr. Parinov using IDT G-Blocks. pGEM LexPR-2A-Cerulean-SV40pA-FRT-Kan-FRT donor plasmid for BAC transgenesis was cloned by Infusion cloning. pGEM BirA-2A-Citrine-SV40pA-FRT-Kan-FRT donor plasmid for BAC transgenesis was cloned by fusion PCR of HA-BirA-2A amplified from PMT-HA-BirA-2A-mCherryRas [Trinh et al.] and insertion into pGEM-GFP-SV40pA-FRT-Kan-FRT where the GFP ORF was been replaced with Citrine. All plasmids generated for the purposes of this study are available from Addgene.

The *TgBAC*(*mpeg1:BirA-Citrine*)*^ox122^* and *TgBAC*(*kita/mitfa:LexPR-Cerulean*)*^ox123/ox124^* transgenic lines was generated by BAC-mediated transgenesis as previously described [Trinh et al.]. Briefly, PCR-amplified BirA-2A-Citrine-SV40pA-FRT-Kan-FRT and LexPR-2A-Cerulean-SV40pA-FRT-Kan-FRT cassettes were recombined into the first coding exon of a driver gene within the corresponding BAC clone (macrophage-specific mpeg1 gene, and melanocyte-specific mitfa or kita, respectively.) In a second recombination step, an iTol2-Ampicillin cassette (provided by Prof Kawakami) was introduced into the BAC backbone as previously published [44, 7]. Wildtype embryos were injected at the one-cell stage with 200 ng/*μ*L of purified BAC DNA and 100 ng/*μ*L tol2 transposase mRNA. Putative founders were outcrossed to wildtype fish and offspring screened for Citrine or Cerulean expression, in combination with PCR amplification of the transgene when expression levels were low. Zebrafish transgenic lines generated and used in this study (Table S1) have been submitted to are available from European Zebrafish Resource Center (EZRC).

### Whole-mount immunofluorescence and imaging

Embryos/larvae were fixed in 4% PFA in Phosphate buffered saline (PBS) (0.1 M, pH 7.4) for 20 minutes at room temperature (RT). Embryos were then rinsed 3 × 15 minutes in PBT (2% DMSO, 0.5% Triton in PBS), incubated in block solution (10% donkey serum in PBT) for 2 hours at RT, followed by incubation with primary antibody (1:200 in block solution) overnight (O/N) at 4 °C. Embryos/larvae were then washed 3-5 times for 1hour at RT, followed by an O/N wash at 4 °C in PBT and then incubated with secondary antibody (1:500 in PBT). Embryos were then washed 6-8 times for 1 hour at RT + O/N at 4 °C in PBT. Primary antibodies used in this study were, mpx (GeneTex, cat #GTX128379) and mpeg1 (GeneTex, cat #GTX54246). The secondary antibody used was AlexaFluor-488-conjugated anti-rabbit IgG (Life technologies, cat #R37116). Citrine signal was amplified using anti-GFP, rabbit polyclonal antibody, AlexaFluor-488 conjugate (Life technologies, cat #A21311) at a 1:500 dilution. mCherry signal was amplified using the RFP-booster that recognises which all dsRed derivatives including mCherry, as per the manufacturers instructions (Chromotek, rba594). Images of stained embryos/larvae were taken on a Zeiss 780 confocal microscope or captured using the Zeiss Z1 lightsheet microscope.

### 5’-ethynyl-2’-deoxyuradine (Edu) labelling of proliferating melanocytes

At 96 hours post fertilization 5 nL Edu (150 *μ*M in 2% DMSO, 0.1% phenol red) was injected into the pericardium of zebrafish embryos. The embryos were incubated overnight, fixed and labelled with the Click-iT Edu AlexaFluor-488 Imaging kit (Invitrogen) as per the manufacturers instructions.

### Tail transection

3 days post fertilisation (dpf) larvae were anaesthetised with 0.4% 3-amino benzoic acidethylester (Tricaine) in E3 and tail transection was performed with a sterile scalpel.

### ’Biotagged’ nuclei Isolation

For nuclei isolation, zebrafish embryos were anaesthetised with 0.01% Tricaine and cranial regions dissected away from anterior yolk sac and Intermediate Cell Mass (ICM). Embryos were washed in hypotonic buffer H (20 mM HEPES (pH 7.9), 15 mM MgCl2, 10 mM KCl, 1 mM DTT, and 1 X Complete protease inhibitor (Roche)) and subsequently re-suspended in 500 *μ*L of buffer H. From here, nuclei isolation was carried out as previously described [11, Trinh et al.]. Following isolation, nuclei-beads were re-suspended in RNA lysis buffer from RNAqueous®- Micro Kit (Life Technologies, cat #AM1931).

### RNA Extraction, library preparation and RNA-seq analysis

Total nuclear RNA was extracted using RNAqueous®-Micro Kit according to manufacturer’s instructions. RNA integrity was checked with a RNA pico chip (Agilent Technologies, cat #5067-1513) using the Agilent 2100 Bioanalyzer. cDNA was synthesized and amplified from 100 pg – 1 ng of input RNA using SMART-seqTMv4 ultra low input kit for RNA (Clontech laboratories, cat # 634888, 634889, 634890, 634891, 634892, 634893, and 634894). Sequencing libraries were prepared using the Nextera XT DNA library preparation kit and Next Generation Sequencing was performed on a NextSeq500 platform using a NextSeq^TM^500 150 cycle High Output Kit) (Illumina, cat #FC-404-1002) to generate 80-basepair paired end reads.

### RNA-seq analysis

Read quality was evaluated using FastQC [Andrews]. Reads were mapped to the (Jul. 2014 Zv10/danRer10 assembly) version of the zebrafish genome using STAR (v.2.4.2a) splice-aware aligner [12]. Count tables were generated using subread FeatureCounts (v1.4.5-p1q), with standard parameters [31]. Differential expression was carried out using DESeq2 R package [33]. Data generated in this study were submitted to GEO (GSE96534)

## Acknowledgements

We thank Isabella Watts for her assistance on the project and Profs. Tudor Fulga, and Ahmed Ahmed for their pertinent comments on the manuscript.

## Author Contributions

AK and TSS conceived and designed the study. AK performed all the experiments. DG carried out bioinformatics analyses and curated the data. AK, DG, GN and TSS analysed the data. AK wrote the first draft of the manuscript. AK, TSS, GN and VC edited the manuscript. TSS supervised the study.

## Data availability

Data generated in this study has been submitted to GEO (GSE96534).

## Supplementary Material

### List of plasmids generated for the purpose of this study available from Addgene

pCrys *β*:ECFP-LexOP:mCherry-NRasK61Q

pCrys *β*:ECFP-LexOP:mCherry-HrasG12V

pCrys *β*:ECFP-LexOP:Cherry-KrasG12V

pGEM LexPR-2A-Cerulean-SV40pA-FRT-Kan-FRT

pGEM BirA-2A-Citrine-SV40pA-FRT-Kan-FRT

Zebrafish transgenic lines used in this study are available from European Zebrafish Resource Center (EZRC) and listed in the Table S1. Two previously generated lines are marked with an asterisk.

**Figure S1.**
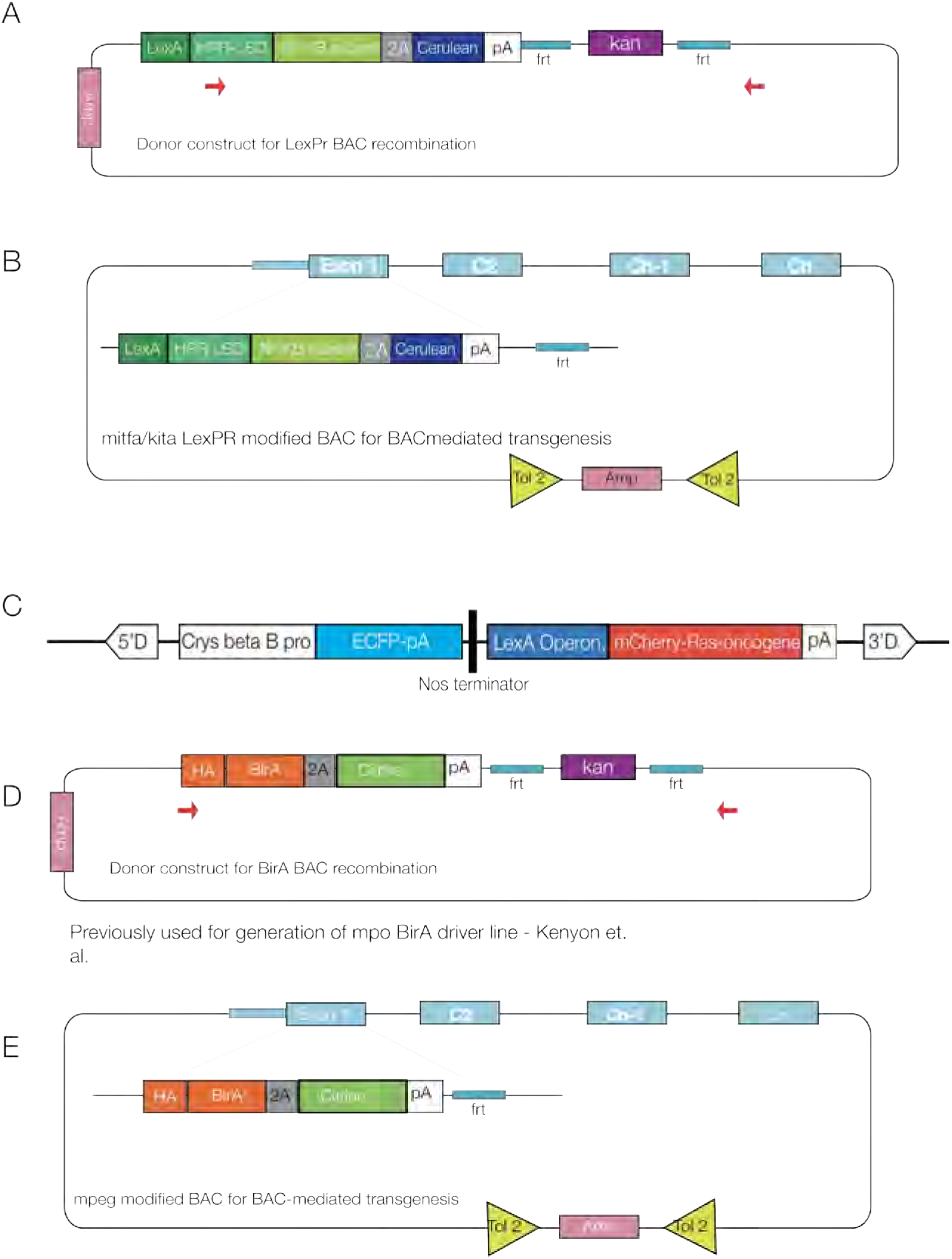
Constructs used in generation of model system. **A)** Schematic of the mpx BAC donor construct, containing HA-tagged BirA (orange), a ribosomal skipping motif - 2A (grey), citrine reporter (green), polyA tail (white), followed by FRT recombination sites (turquoise) flanking a kanamycin selection cassette. Ampicillin selection cassette, not amplified as a part of recombination cassette, is used as a selection marker for E. coli during plasmid DNA isolation. Red arrows indicate position of primers used for amplification and recombination into the mpx BAC. **(B)** Schematic of kita/mitfa modified BAC DNA containing the LexPR transactivator (green), a viral self-cleaving peptide (2A, grey) and a fluorescent reporter (cerulean, blue), followed by a polyA tail (pA, white) and remaining Frt site, recombined into the first exon, with an ampicillin selection cassette (Amp, pink) and tol2 arms (yellow) on the BAC plasmid backbone for BAC mediated transgenesis. **(C)** Schematic PCrysB:ECFP-LexOP- mCherry-RasOncogene construct, embedded within a non-autonomous Ds element to produce Ds insertions in the zebrafish genome with the aid of a modified Ac transposase.**(D)** Schematic of the mpeg1 BAC donor construct, containing HA-tagged BirA (orange), a ribosomal skipping motif - 2A (grey), citrine reporter (green), polyA tail (white), followed by FRT recombination sites (turquoise) flanking a kanamycin selection cassette. Ampicillin selection cassette, not amplified as a part of recombination cassette, is used as a selection marker for E. coli during plasmid DNA isolation. Red arrows indicate position of primers used for amplification and recombination into the mpeg1 BAC. **(E)** Schematic of mpeg1 modified BAC DNA with HA-tagged BirA (orange), a ribosomal skipping motif - 2A (grey), citrine reporter (green), polyA tail (white), followed by the remaining FRT sites (turquoise) recombined into the first exon, with a BAC-specific ampicillin-Tol2 cassette (iTol2) in pink and yellow.

**Figure S2.**
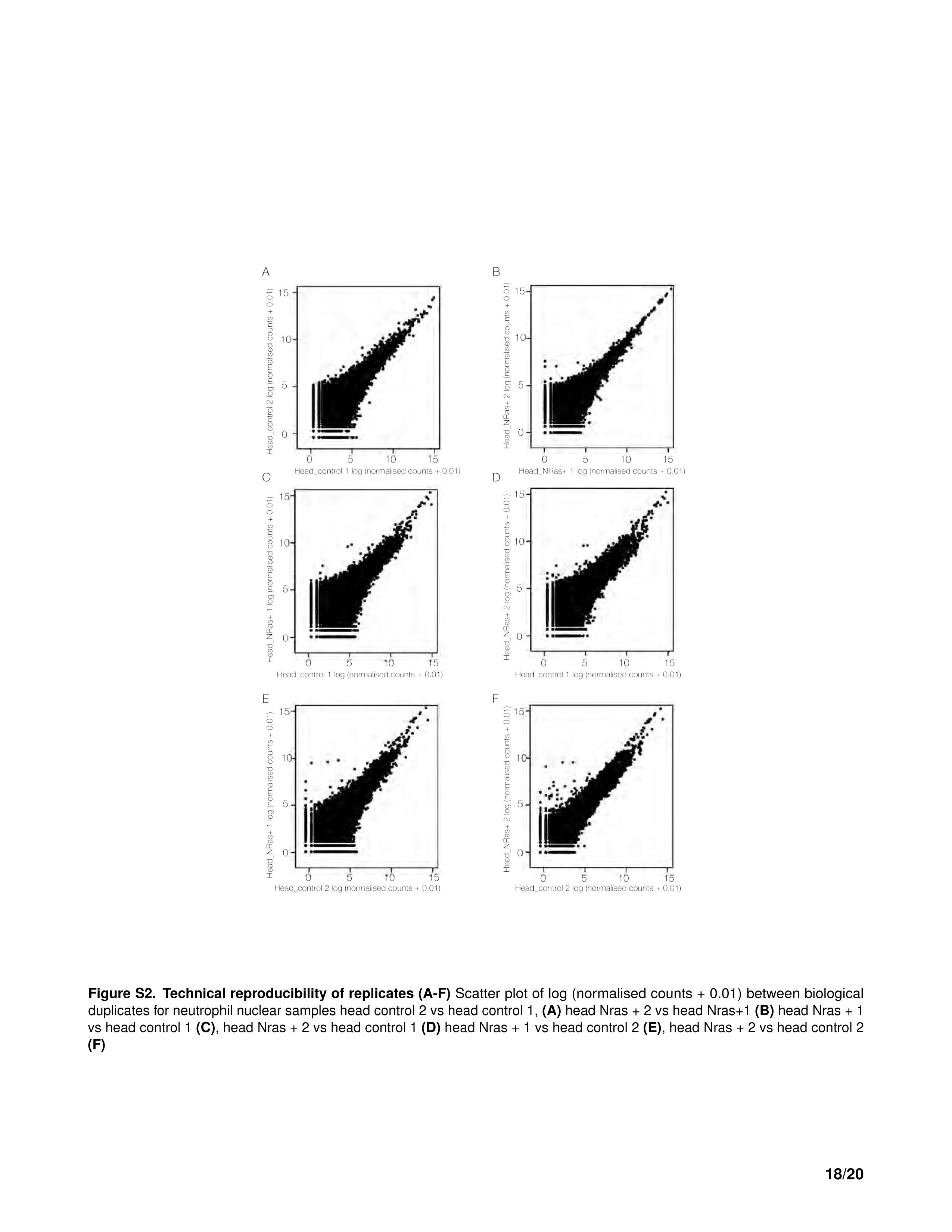
Technical reproducibility of replicates (A-F) Scatter plot of log (normalised counts + 0.01) between biological duplicates for neutrophil nuclear samples head control 2 vs head control 1, **(A)** head Nras + 2 vs head Nras+1 **(B)** head Nras + 1 vs head control 1 **(C)**, head Nras + 2 vs head control 1 **(D)** head Nras + 1 vs head control 2 (**E**), head Nras + 2 vs head control 2 (**F**)

**Figure S3.**
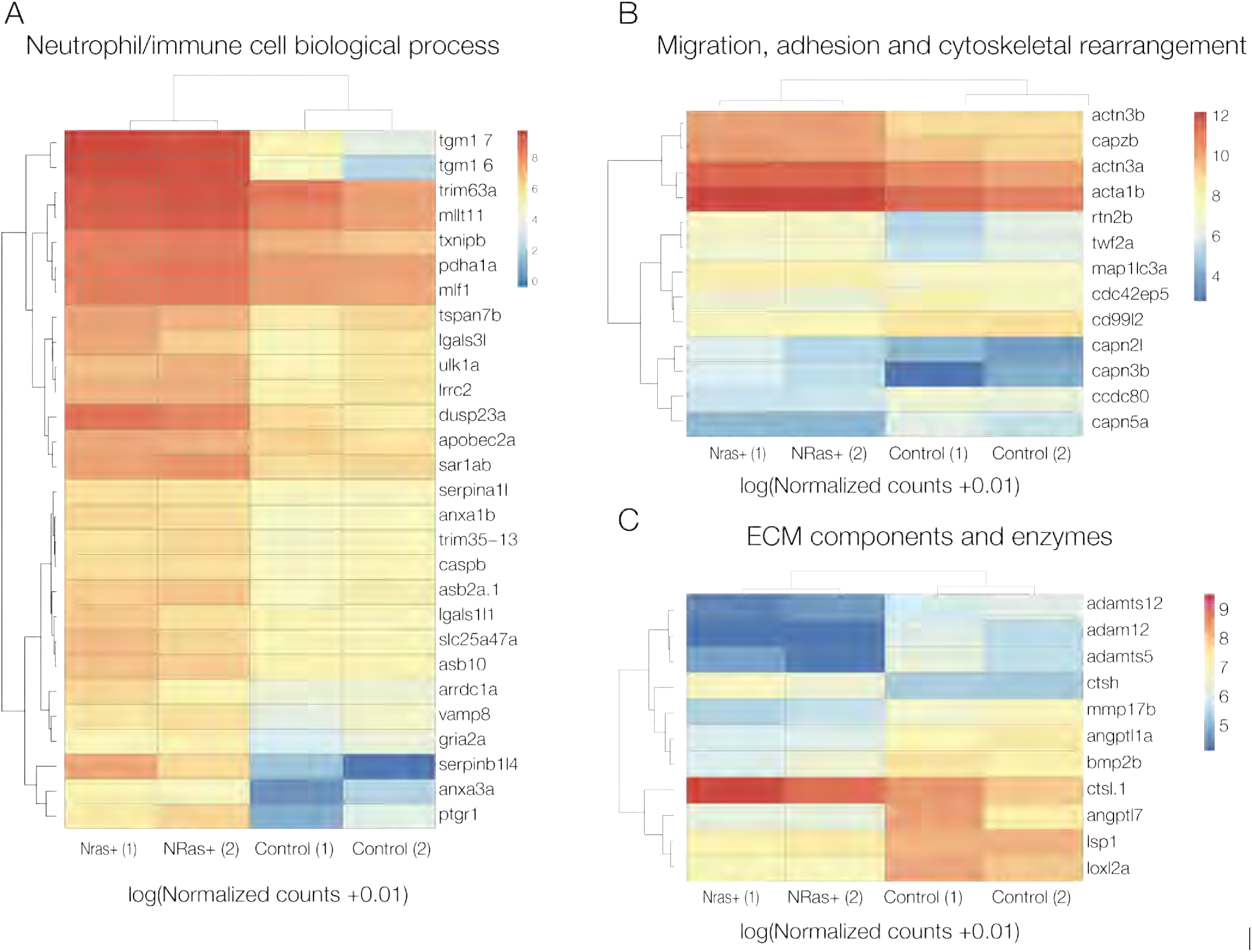
Classification of differentially expressed transcripts (A-C) Heatmaps show the log10 (normalised counts (NMCT) +0.01) of selected differentially expressed transcripts (adjusted p-value ί 0.05). Red - high expression. Yellow - medium expression. Blue - low expression. Neutrophil/immune cell biological processes (A), Migration, adhesion and cytoskeletal rearrangement **(B)** and ECM components and enzymes **(C)**

**Table S1:**
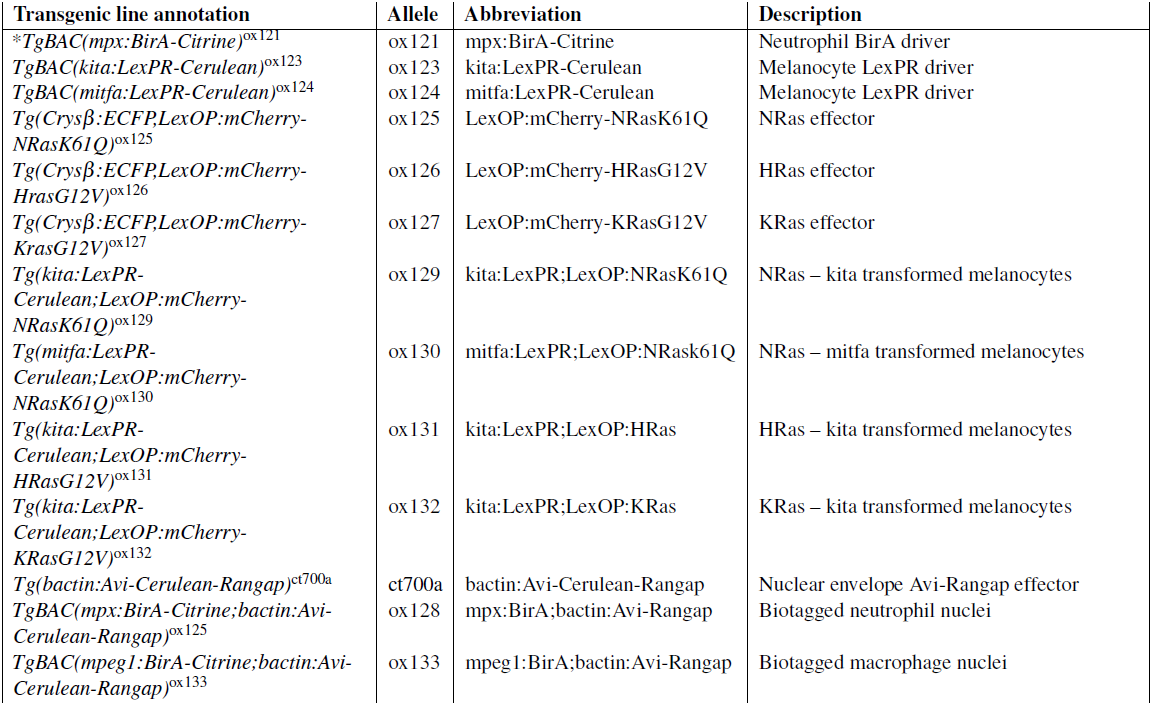

